# An mRNA vaccine against SARS-CoV-2: Lyophilized, liposome-based vaccine candidate EG-COVID induces high levels of virus neutralizing antibodies

**DOI:** 10.1101/2021.03.22.436375

**Authors:** H. Christian Hong, Kwang Sung Kim, Shin Ae Park, Min Jeong Chun, Eun Young Hong, Seung Won Chung, Hyun Jong Kim, Byeong Gyu Shin, Abdennour Braka, Jayaraman Thanappan, Sunghoon Jang, Sangwwok Wu, Yang Je Cho, Seok-Hyun Kim

## Abstract

In addition to the traditional method of vaccine development, the mRNA coronavirus vaccine, which is attractive as a challenging vaccination, recently opened a new era in vaccinology. Here we describe the EG-COVID which is a novel liposome-based mRNA candidate vaccine that encodes the spike (S) protein of SARS-CoV-2 with 2P-3Q substitution in European variant. We developed the mRNA vaccine platform that can be lyophilized using liposome-based technology. Intramuscular injection of the EG-COVID elicited robust humoral and cellular immune response to SARS-CoV-2. Furthermore, sera obtained from mice successfully inhibited SARS-CoV-2 viral infection into Vero cells. We developed EG-COVID and found it to be effective based on *in vitro* data, and we plan to initiate a clinical trial soon. Since EG-COVID is a lyophilized mRNA vaccine that is convenient for transportation and storage, accessibility to vaccines will be significantly improved.

## Introduction

COVID-19 is caused by the coronavirus, which is an enveloped RNA virus that is widely distributed in humans, other mammals, and birds and causes respiratory, intestinal, liver, and nerve diseases. (Weiss et al., 2011). In addition to the very common coronavirus responsible for the human common cold, there are three coronaviruses that cause serious illness in humans: SARS-CoV, MERS-CoV, and SARS-CoV-2 (Cox et al., 2020). In December 2019, an outbreak of severe acute respiratory syndrome coronavirus 2 (now namely, SARS-CoV-2) infection occurred at seafood market in Wuhan, Hubei Province, China, and spread across China (Zhu et al., 2020). Considering the rapid spread of this virus and its impact on an international scale, COVID-19 was declared a pandemic by the World Health Organization on March 11, 2020 (WHO, 1 DEC 2020). The urgent development of safe and effective vaccines in the global coronavirus pandemic has become a common goal for humanity, where numerous multinational pharmaceutical companies and biotechnology companies are immersed in the development of COVID-19 vaccines around the world simultaneously.

The mRNA-based vaccines are in the limelight because of their potential for rapid development than the conventional vaccines at the COVID-19 pandemics. The history of mRNA-based vaccines is relatively short, and its first application was introduced in the 1990s, using mice to intramuscularly inject mRNA to locally produce an encoded reporter protein (Wolff et a.,1990). The mRNA vaccine combines the benefits of the subunit vaccine and the live attenuated vaccine, and there is no risk associated with the live attenuated vaccine or the DNA vaccine (Reichmuth et al., 2016). Of all the viral antigens studied, the spike (S) and nucleocapsid (N) proteins are considered to be important immunogenic antigens for SARS-CoV-2 (Assadiasl, 2020). Among these, most vaccine candidates use the coronavirus glycosylated spike protein (S) as the primary antigenic target.

Liposomes are defined as phospholipid vesicles consisting of one or more concentric lipid bilayers enclosing discrete aqueous spaces and its use as pharmaceutical applications are well known (Ickenstein et al., 2019, Torchilin, 2005, Sercombe et al., 2015). In 1970, liposomes were proposed as a drug carrier for altering the therapeutic index of a drug by reducing toxicity or increasing the efficacy of the parent drug (Lian et al., 2001). Vartak and colleagues suggested the use of liposome as antigen or immunomodulatory molecules carrier (Vartak et al., 2016). Furthermore, cationic liposome is considered as an effective adjuvant for vaccine delivery as well as enhancing both cellular and humoral immune reactions (Mai et al., 2020). As a reference, Herpes zoster vaccines are reported to use neutral or cationic liposome as antigen delivery systems (Lal et al., 2015, Cunningham et al., 2016, Wui et al, 2019, and 2021). Cationic liposomes have many important safety advantages, especially when making viral vectors, are less expensive, have no restrictions on the size of nucleic acids, and are easily accessible without special experience in their use and handling (Karmali et al., 2007). Since the cationic liposome is positively charged, it is able to hold nucleic acids such as mRNA that is negatively charged and deliver mRNAs to cells and tissues efficiently. When prepared under proper conditions, these lipoplexes (lipid-nucleic acid complex) retain their full positive charge, allowing them to efficiently bind to negatively charged cell membranes, enter the cells, protects nucleic acids from nucleases in the serum or cytoplasm (Karmail et al., 2007).

By the way, all currently developed mRNA-based vaccines employ the lipid nanoparticles (LNP) as their delivery vehicles for mRNA. LNPs are composed of ionizable lipid, helper lipid, cholesterol, and PEG-lipid (Kim, et al., 2021). Although LNPs are very efficient tools for delivery of mRNA into human body, all the LNP based vaccines are distributed in liquid form only, and require special cold chain system to maintain their efficacy.

In this report, we established cationic liposome based mRNA delivery system and developed a lyophilizable SARS-CoV2 candidate vaccine (EG-COVID) with mRNA encoding the full spike glycoprotein (S) of the European mutant strain (D614G) with 2P-3Q substitution (Bangaru et al., 2020). We found that EG-COVID could elicit robust humoral and cellular immune response to SARS-CoV-2, furthermore, sera obtained from mice successfully inhibited SARS-CoV-2 viral infection into Vero cells. So here we want to describe the immunogenicity and neutralizing antibody activity of the EG-COVID.

## Author’s contributions

1. Conception or design of the work: Y.J. C., S-H.K., K.S.K., H.C.H., A.B., J.T., S.J., S.W.
2. Data collection: Y.U.H., M.J.C., S.A.P.
3. Data analysis and interpretation: Y.U.H., M.J.C., S.A.P., S.W.C., H.J.K., B.G.S.
4. Writing the article: H.C.H., S-H. K.
5. Critical revision of the article: Y.J. C., S-H. K.
6. Funding acquisition: Y.J.C
7. Final approval of the version to be published: All authors reviewed and approved the final article.

## Competing interests

The authors declare no conflict of interest. Corresponding authors:

Yang Je Cho and/or Seok-Hyun Kim

## Acknowledgement

We thank Doo Sik Kim and Young Hee Cho for proofreading the manuscript.

## Results

### Design and optimization of CoV2-F004 (SARS-CoV-2-2P-3Q full-length spike)

It was published that the substitution of K986P and K987P (2P) increase stability of the protein, and the substitution of Prolines to Glutamate in S1/S2 polybasic cleavage site (3Q) confers the protease resistance (Corbett et al., 2020, Torchilin, 2005). Based on these findings, we used the mRNA sequence encoding the SARS-CoV-2 Spike protein with 2P-3Q substitutions in European variant (D614G) and named CoV2-F004. The translated Spike protein from CoV2-F004 was predicted to be superior to Wuhan variant (614D) in structural stability and antigen presentation ability through AI-modeling simulation.

### In vitro transcription of CoV2-F004 mRNA

CoV2-F004 mRNA was produced by TriLink biotechnologies with Cleancap® technology to increase translation efficiency. 5-Methoxyuridine was used to reduce immune response of cells against mRNA. Produced mRNA demonstrated an expected size of single band in agarose gel electrophoresis (Figure 1C), demonstrating the functionality of CoV-2-F004 structure.

**Figure 1.**
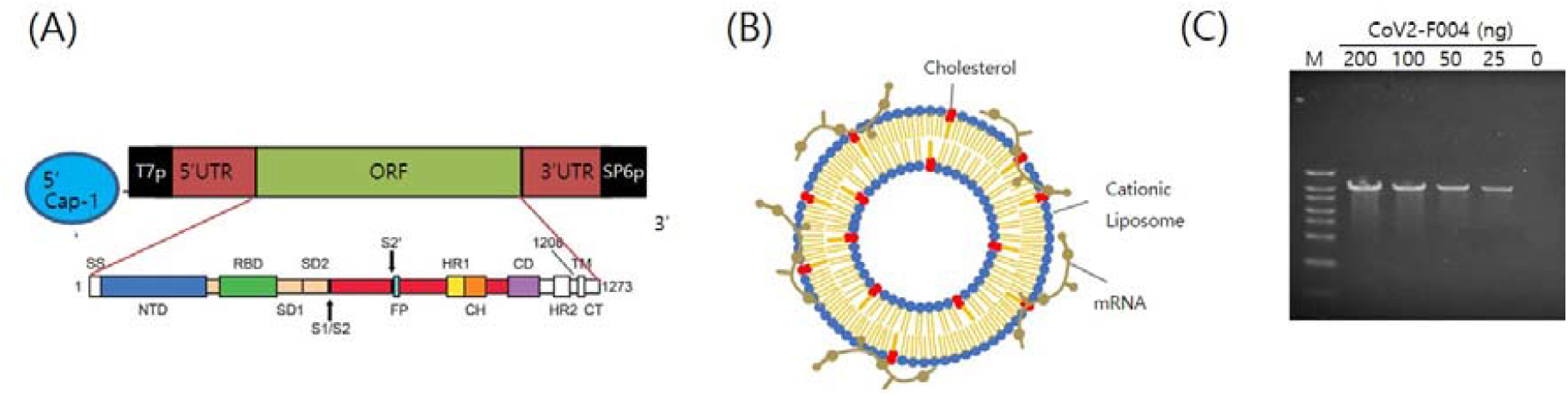
The structure of EG-COVID vaccine. (A) A schematic illustration of antigen coding mRNA Cov2-F004 structure. (B) Schematic illustration of the structure of EG-COVID. (C) The functionality of CoV2-F004 Structure. In vitro transcription of CoV2-F004 showed the increase of transcript according to CoV2-F004 amount used in the transcription. M; size marker.

### Characterization of EG-COVID

Liquid form of EG-COVID (L-EG-COVID) and reconstituted lyophilized form of EG-COVID (F-EG-COVID) were analyzed for hydrodynamic size and zeta potential. L-EG-COVID and F-EG-COVID had a Z-average diameter of 191.7 ± 8.5 nm and 266.7±12.2 nm with a polydispersity index (PDI) of 0.2 and a zeta potential of –54.5 ± 3 mV and –44.4 ± 2.1 mV, respectively (Figure 2A, 2B, 2D and 2E). Cryo-transmission electron microscopy (Cryo-TEM) of L-EG-COVID and F-EG-COVID revealed that both of EG-COVID mostly forms small unilamellar vesicles (SUV) with diameter of 80-100 nm (Figure 2C and Figure 2F).

**Fig 2.**
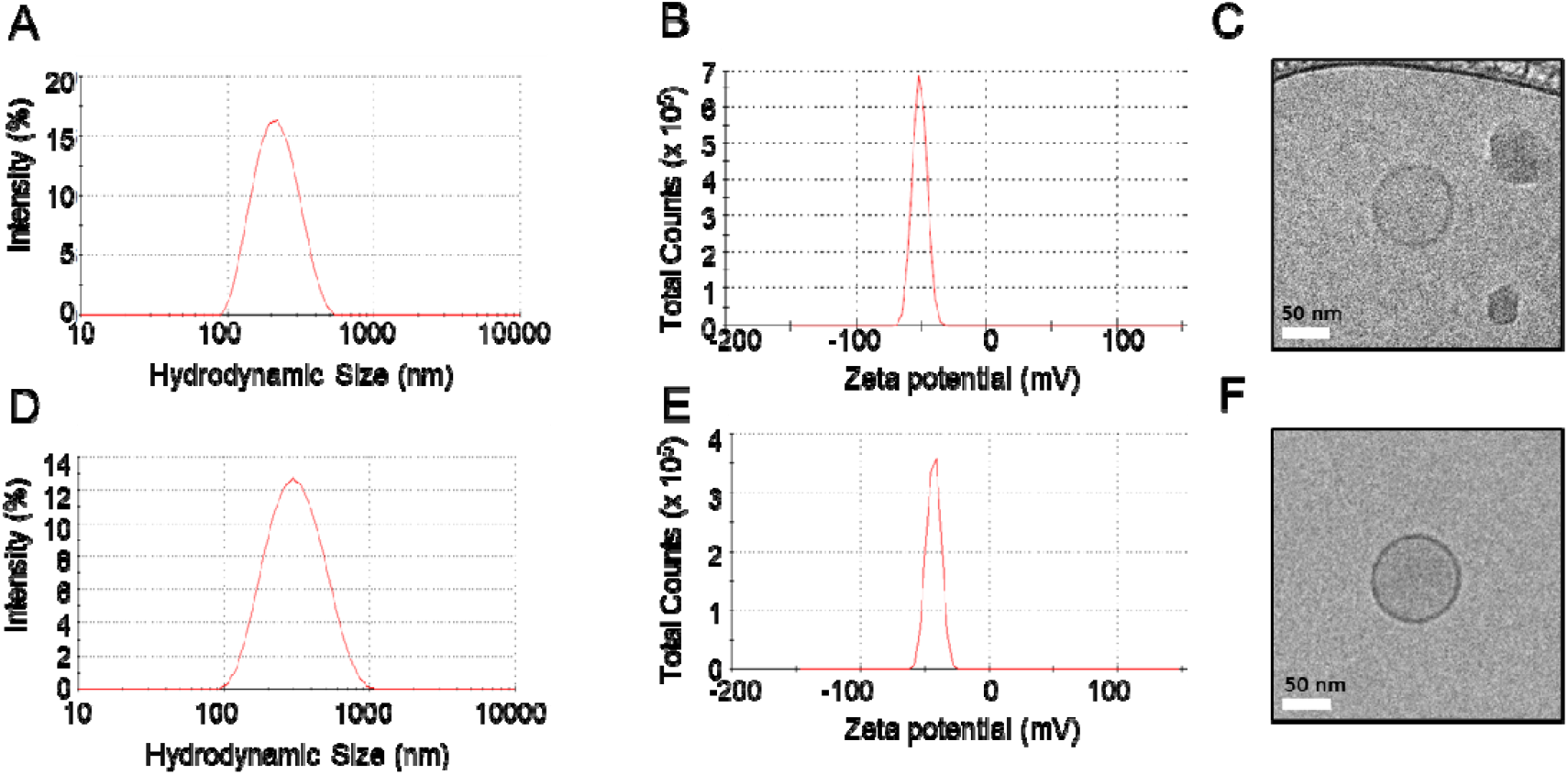
Physical characteristics of the EG-COVID. Representative DLS size distribution (A), zeta-potential (B), and Cryo-TEM image (C) of EG-COVID; Representative DLS size distribution (D), zeta-potential (E), and Cryo-TEM image (F) of F-EG-COVID. Scale bar □= □ 50□nm.

### Cationic liposomes as mRNA delivery carriers

In order to confirm the ability of cationic liposomes as mRNA delivery carriers, complexes of cationic liposomes and *Renilla* luciferase mRNA were prepared and intramuscularly injected into mice to confirm the mRNA delivery ability of cationic liposomes. We used 1 to 20 μg of mRNA complexed with cationic liposomes and observed that the expression of *Renilla* luciferase was successfully achieved. The level of expression of 10 ug of mRNA with cationic liposome was similar to the level of 10 μg of mRNA delivered using lipid nano particle. Therefore, we concluded that our liposome complex is nicely functional as mRNA delivery carriers for mRNA vaccines (Figure 3A). In addition, we prepared a lyophilized form of the *Renilla* luciferase mRNA-cationic liposome complex and confirmed that the lyophilized form of the complex has similar mRNA delivery capability to that of the liquid form (Figure 3B).

**Figure 3.**
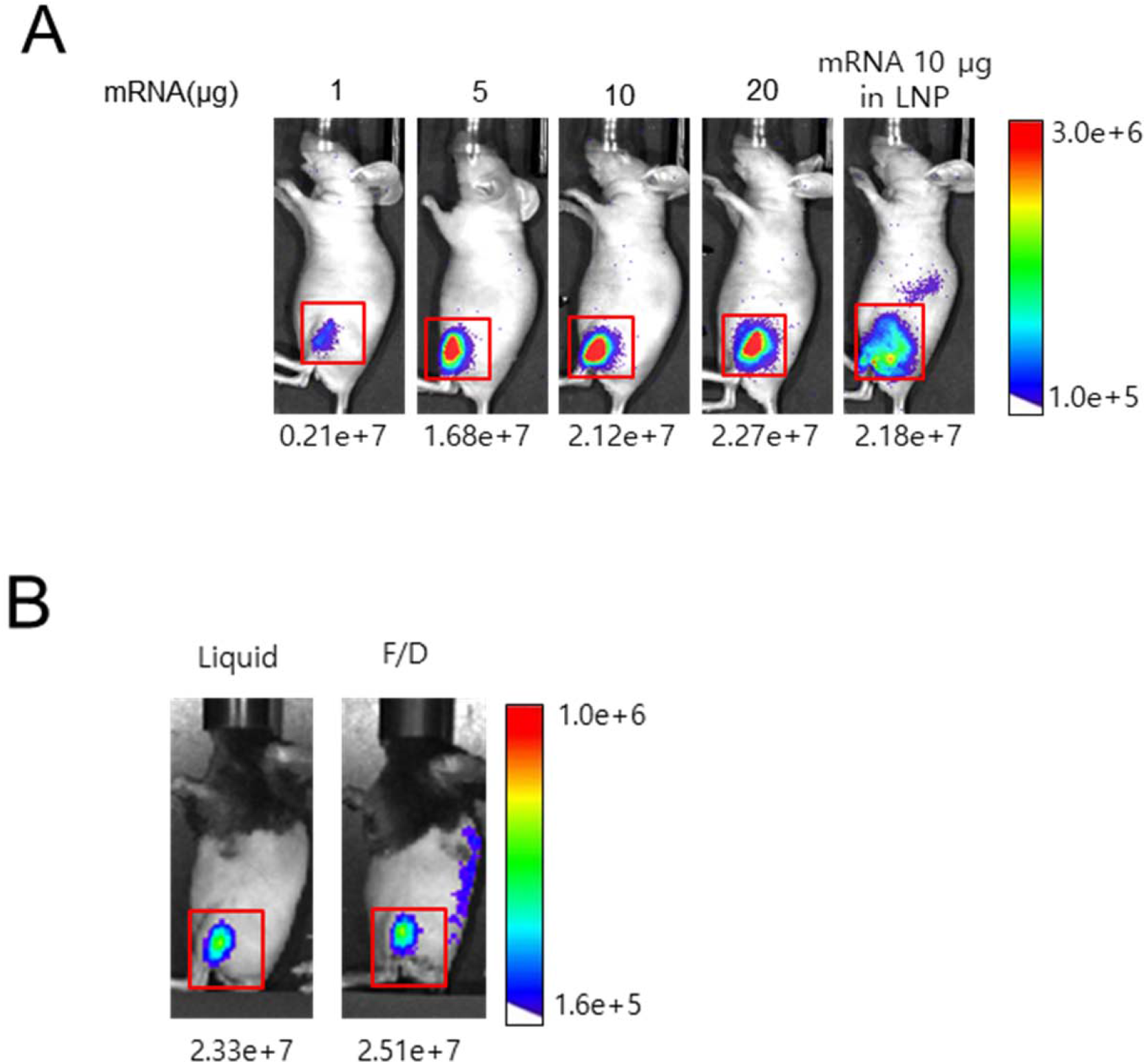
Cationic Liposomes as Efficient mRNA Carriers. (A) 1, 5, 10, and 20 μg of Renilla luciferase mRNAs were complexed with cationic liposomes and intramuscularly injected into mice and bioluminescence was measured for 60 seconds after administration of Renilla luciferase substrate via tail vein. 10 μg of Renilla luciferase mRNA complexed in Lipid nano particle was used as a control (B) Liquid and lyophilized (F/D) forms of Renilla luciferase mRNA-cationic liposomes were injected and bioluminescence was measured. 10 μg of Renilla luciferase mRNA was used in each form.

### In vitro Expression of EG-COVID

We have generated a complex with cationic liposomes as a delivery system of Cov2-F004 naming them as EG-COVID and verified the ability to deliver Cov2-F004 mRNA and expression of SARS-CoV2 spike protein *in vitro* (Figure 4). Re-hydrated lyophilized form of EG-COVID 001, 002, 003 and 004, which were prepared differently in their lipid composition, were treated in the culture medium of 293T cells, and expression of SARS-Cov2 protein was confirmed by western blot after 24-hour incubation. It was confirmed that there was some difference in the expression of protein depending on the lipid composition of EG-COVID.

**Figure 4.**
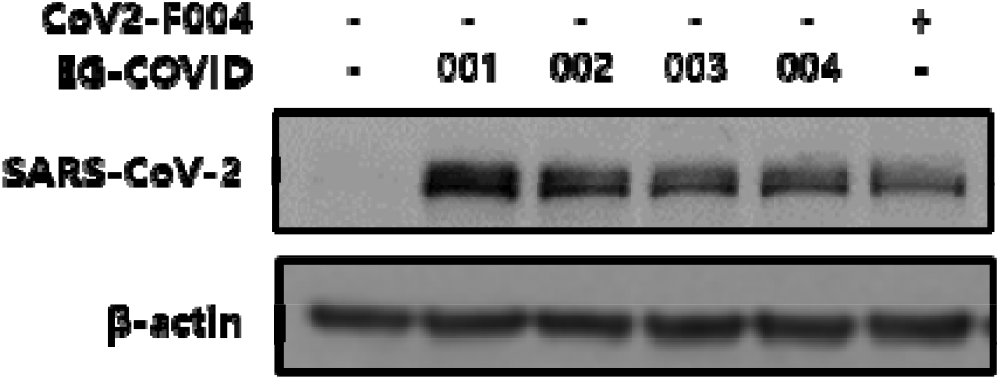
Expression of SARS-CoV-2 antigen by EG-COVID. Four EG-COVID formulations generated with different lipid compositions were introduced into 293T cells and the expression of CoV2-F004 was confirmed by western blot using antibody against SARS-CoV2 protein. All EG-COVID formulations were re-hydrated from lyophilized forms. CoV2-F004 mRNA alone was introduced to 293T cells using Lipofectamine messengerMax RNA kit and used as a control.

### Immunogenicity of EG-COVID

Female B6C3F1/slc mice were immunized by intramuscular administration of L-EG-COVID and F-EG-COVID twice at 3-week interval. After 2 weeks of last administration, mice were sacrificed to evaluate immunogenicity of EG-COVID. We used both liquid (L-EG-COVID) and lyophilized (F-EG-COVID) form of EG-COVID in the experiment.

Humoral immunity was confirmed by anti-RBG antibody titer, and cellular immunity was confirmed by the amount of IFN-γ secreted from splenocytes stimulated with S1 and S2 peptides. We found that both liquid and lyophilized form of EG-COVID induced humoral and cellular immunity successfully (Figure 5A and 5B).

**Figure 5.**
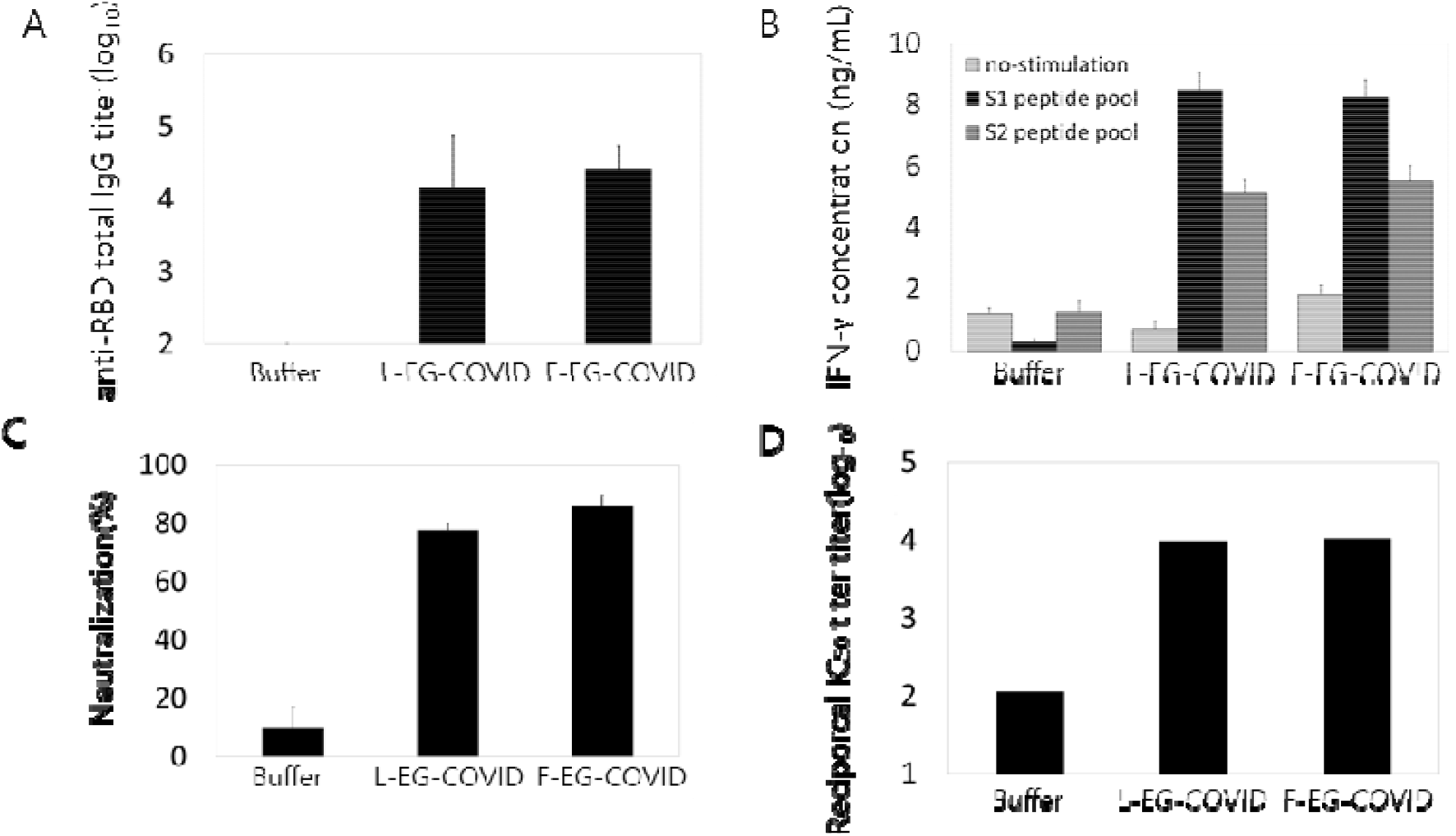
Immunogenicity of EG-COVID and neutralizing antibody reaction induced by EG-COVID immunization. (A) Anti-RBD-specific total IgG antibody titers in sera were measured by an end-point dilution ELISA method. (B) Amount of IFN-_γ_ secreted into the culture medium from stimulated splenocytes were measured by sandwich ELISA method after 72 hours. (C) sVNT (surrogate virus neutralization test) was used to assess the neutralizing ability of the RBD protein (p <0.001, compared with the buffer by Student’s t test). (D) Viral neutralizing activity of immunized sera was measured by PRNT (plaque reduction neutralization test) and IC_50_ was calculated.

We also verified that the antibody formed by EG-COVID actually has neutralizing ability against SARS-CoV2 virus. The result obtained using surrogate virus neutralization tests which assess RBD binding ability of antibodies showed that EG-COVID immunization successfully generated RBD binding antibodies in serum. Furthermore, EG-COVID immunized serum significantly inhibited plaque formation induced by Wuhan variant of SARS-CoV2 virus infection into VERO cells. There was no difference in EG-COVID immunization ability between liquid and lyophilized formulations (Figure 5C and 5D). Therefore, we can conclude that EG-COVID induced a sufficient level of humoral and cellular immunity against SARS-CoV-2 virus and is a first-class candidate for mRNA vaccine that can be easily lyophilized.

## Materials and Methods

### Lipids and chemicals

1,2-dioleoyl-3-trimethylammonium-propane (DOTAP) and Cholesterol were purchased from Merck Millipore (Darmstadt, Germany) or Avanti Polar lipid (AL, USA). 1,2-dioleoyl-sn-glycero-3-phosphoethanolamine (DOPE) and 1,1’-((2-(4-(2-((2-(bis(2-hydroxydodecyl)amino)ethyl)(2-hydroxydodecyl)amino)ethyl)piperazin-1-yl)ethyl)azanediyl)bis(dodecan-2-ol) (C12-200) were from Corden Pharma (Plankstadt, Germany). 1,2-distearoyl-sn-glycero-3-phosphocholine (DSPC) and 1,2-dimyristoyl-rac-glycero-3-methoxypolyethylene glycol-2000 (DMG-PEG2000) were obtained from Avanti Polar Lipid (AL, USA). Hydroxyethyl piperazine Ethane Sulfonic acid (HEPES) solution was purchased from Sigma Aldrich (MO, USA). Sucrose was obtained from Merck Millipore (Darmstadt, Germany). 1 M Sodium acetate solution (pH 4.6) were from T&I (Gangwon-do, Korea).

### mRNAs

*Renilla* luciferase mRNA was purchased from TriLink (CA, USA). CoV2-F004 mRNA was produced by TriLink (CA, USA)

### Preparation of EG-COVID and mRNA-encapsulated LNP

Cationic liposomes were prepared with 1,2-dioleoyl-3-trimethylammonium-propane (DOTAP; Merck Millipore, Darmstadt, Germany), 1,2-dioleoyl-sn-glycero-3-phosphoethanolamine (DOPE; Corden Pharma, Plankstadt Germany) and Cholesterol (Merck Millipore, Germany) by a thin-film method modified from previously described by Wui (Wui et al., 2019). Briefly, for the thin-film method, a chloroform solution containing the DOTAP: DOPE: Cholesterol mixture was dried in a round bottom flask using a rotary evaporator (IKA, Germany) to form a thin-lipid film. The lipid film was rehydrated with a nuclease free-4% (w/v) sucrose solution (Sigma Aldrich, MO, USA) in 20 mM HEPES (pH7.4; Sigma Aldrich, MO, USA) and homogenized using a microfluidizer (Avestin, ON, Canada). The mRNA CoV2-F004 were coated to the cationic liposome. The EG-COVID was lyophilized using a freeze-dryer (Lyoph-Pride, Ilshin BioBase, Gyeonggi-do, Korea).

LNP were prepared using microfluidic mixer Ignite™ (Precision NanoSystems, Vancouver, BC, Canada) as previously described by Sabnis and colleagues (Sabnis. et al. 2018) In brief, 1,1’-((2-(4-(2-((2-(bis(2-hydroxydodecyl)amino)ethyl)(2-hydroxydodecyl)amino)ethyl)piperazin-1-yl)ethyl)azanediyl)bis(dodecan-2-ol) (C12-200; Coden Pharma, Germany), 1,2-distearoyl-sn-glycero-3-phosphocholine (DSPC; Avanti Polar Lipid, AL, USA), Cholesterol (Merck Millipore, Germany) and 1,2-dimyristoyl-rac-glycero-3-methoxypolyethylene glycol-2000 (DMG-PEG2000; Avanti Polar Lipid, AL, USA) were resolved in ethanol at molar ratios of 50:10:38.5:1.5. The lipid mixture was obtained with 25 mM sodium acetate buffer (pH 3; T&I, Gangwon-do, Korea) containing *Renilla* luciferase mRNA (TriLink, CA, USA) at a N/P ratio of 5.6. using microfluidic mixer Ignite™. Formulations were dialyzed against nuclease free-4% (w/v) sucrose solution in 20 mM HEPES (pH7.4). The EG-COVID and mRNA-encapsulated LNP were stored at 4°C and used within a week.

### Measurement of particle size and zeta potential of EG-COVID

The particle size and polydispersity index (PDI) of liposomes were determined by dynamic light scattering (DLS) using a Zetasizer Nano ZSP (Malvern Instruments, Worcestershire, UK). Malvern Zetasizer DTS software (version 7.12) was used for data acquisition and analysis. All measurements were carried out in triplicate.

### Morphology of EG-COVID

The cryogenic transmission electron microscopy (Cryo-TEM) was used for determination of morphology of EG-COVID. Liquid form of EG-COVID or lyophilized EG-COVID powder was rehydrated with deionized water were used for Cryo-TEM analysis. Three microliters of the liposome preparation were placed on a 200 mesh copper grid of lace carbon film and semi-automatically vitrified using a Vitrobot ™ (FEI, OR, USA). Vitrified samples were maintained at liquid nitrogen temperatures during sample transfer and imaged with a Tecnai F20 G2 (FEI, OR, USA) at 120 keV.

### Cell culture and mRNA transfection

HEK-293T cell was obtained from ATCC (Manasass, VA, USA). Cell was cultured in Dulbecco’s modified Eagle media (DMEM) (Corning, NY, USA) containing 1x penicillin/streptomycin (Thermo Scientific, MA, USA) and 10 % fetal bovine serum (FBS) (Hyclone, Marlborough, IL, USA) in a humidified atmosphere containing 5% CO2 at 37 °C. Transfection of CoV2-F004 mRNA was performed using Lipofectamin messengerMax RNA (Invitrogen, thermo Scientific, MA, USA) in accordance with the manufacturer’s instructions. EG-COVID 001, 002, 003 and 004 (10 ug mRNA containg complex each) were introduced into 293T cells by 18 hour incubation in OPTi-MEM (Thermo Scientific, MA, USA).

### Western blotting

Cells were washed twice in phosphate-buffered saline (PBS) (WELGENE, Kyunggi-do, Korea) and lysed in lysis buffer (Cell signaling, Danvers, MA, USA) with protease inhibitor (Cell signaling, Danvers, MA, USA). Lysates were cleared by centrifugation and supernatants were obtained. Equal amounts of proteins were resolved on 8% Bis-Tris page gel (Invitrogen, Thermo Scientific, Waltham, MA, USA). Proteins were resolved eletro-trasferred onto polyvinylidene fluoride (PVDF) membranes (GE Healthcare, Chicago, IL, USA). The membranes incubated with the appropriate antibody in Tris buffered with Tween-20 (TTBS). The primary and secondary antibodies were SARS-CoV-2-S1 (Novus, Centennial, CO, USA), beta-actin (Cell signaling, Danvers, MA, USA) and anti-rabbit (Invitrogen, Thermo Scientific, Waltham, MA, USA).

### Bioluminescence imaging

For bioluminescence imaging studies, 6-week-old female Balb/c nude or C57BL/6N mice (SLC, Japan) were used. The mice received *Renilla* luciferase mRNA-liposome (or lipid nanoparticle) complex intramuscularly. 6 hours after the administration, mice were anesthetized with intraperitoneal injection of Avertin working solution (Sigma Aldrich, MO, USA), then, 0.15 mg/mL ViviRen luciferase substrate (Promega, WI, USA) were injected intravenously as 1mg/kg. The bioluminescence imaging was obtained with Ami HTX (Spectral Instruments Imaging, AZ, USA) immediately after the injection of substrate. All images were taken under 60 seconds of exposure time and analyzed with Aura software (Spectral Instruments Imaging, AZ, USA). The regions of interest (ROIs, photons/s) were manually selected based on the signal intensity.

### Immunization of mice

For immunogenicity studies, 6-week-old female B6C3F1 mice (SLC, Japan) were randomly assigned into experimental groups (n = 6). The Mice were immunized intramuscularly two doses at 3-week intervals with an EG-COVID (L-EG-COVID, F-EG-COVID). Two weeks after the final immunization, blood was obtained by cardiac puncture and spleen were isolated from the immunized mice. Sera were obtained by centrifugation at 3,000 RPM for 5 minutes at room temperature and kept at −70°C deep freezer. After extracting the spleen from the immunized mice, the spleens of each group were pooled and washed in PBS (T&I, Gangwon-do, Korea) with 1% Penicillin/Streptomycin (Gibco, MA, USA). A 70 μm cell strainer (VWR life science, PA, USA) was used to homogenize the tissue. Homogenized tissues were transferred to a 15 mL tube (SPL, Gyeonggi-do, Korea), followed by centrifugation at 4° C, 3,000 rpm for 5 minutes, and then the supernatant was removed. The cells were suspended in 3 mL of RBC lysis buffer (Biolegend, CA, USA), allowed to stand at room temperature for 3 minutes, and then centrifuged at 4° C, 3,000 rpm for 5 minutes, and then the supernatant was removed. Cells were suspended by adding 3 mL of PBS with 1% Penicillin/Streptomycin, followed by centrifugation at 4° C, 3,000 rpm for 5 minutes to wash the cells. Discard the supernatant and RPMI1640 (Gibco, MA, USA) [10% FBS (Hyclone, MA, USA), 100 μ/mL penicillin (Gibco, MA, USA), 100 μg/mL streptomycin (Gibco, MA, USA), 0.5 ng/mL recombinant mouse IL-2 (R&D system, MN, USA), and 50 μM β-mercaptoethanol (Sigma Aldrich, MO, USA)] was added at 2 × 10^7^ cells/mL.

### Enzyme-linked Immunosorbent Assay (ELISA)

Antigen-specific antibody titer was determined by end-point dilution ELISA. A 96 well immunoplate (Thermo Fisher, MA, USA) was coated with 100 ng/well of SARS-CoV-2 RBD protein (mybiosource, CA, USA) for 16 hours at 4°C. The plates were blocked with 1% bovine serum albumin (MP Biomedicals, CA, USA) in phosphate buffered saline (PBS) for 1hr at 37 °C, and incubated with sera that was serial diluted with 1% bovine serum albumin in PBS for 1 hour at 37°C. Bound antibodies were detected with HRP-conjugated goat anti-mouse total IgG antibody (Jackson, PA, USA) diluted 1:5000 for 1 hour at 37°C. The color reaction was developed by adding KPL sureblue TMB (3,3’,5,5’-tetramethylbenzidine) microwell peroxidase substrate (Seracare, MA, USA) and 1N Sulfuric acid (Daejung, Busan, Korea) was added to stop the reaction. Absorbance was measured on an Epoch Microplate Spectrophotometer (Biotek, VT, USA) at 450 nm (Ryu, et al., 2016).

### In vitro re-stimulation and cytokine analysis

Splenocytes were resuspended in RPMI1640 (Gibco, MA, USA) supplemented with 10% FBS (Hyclone, MA, USA), 100 u/mL penicillin (Gibco, MA, USA),100 μg/mL streptomycin (Gibco, MA, USA), 0.5 ng/mL recombinant mouse IL-2(R&D system, MN, USA), and 50 μM β-mercaptoethanol (Sigma Aldrich, MO, USA). Cells were seeded at 2 × 10^6^ cells/well in a 96 well culture plate (SPL, Gyeonggi-do, Korea) and stimulated with two pools of SARS-CoV-2 spike peptide (JPT, Germany) or media. Each peptide pool (JPT1, S1 peptide pool; JPT2, S2 peptide pool) was used at 1 μg/mL (Corbett, et al, 2020). After 72 hours of incubation at 37°C, culture media were collected for assay. IFN-γ secreted from splenocytes were measured using mouse IFN-γ Duoset ELISA kit (R&D system, MN, USA).

### Surrogate Virus Neutralization Test (sVNT)

The surrogate virus neutralization test (sVNT) assay was performed using the SARS-CoV-2 surrogate virus neutralization test kit (GenScript, NJ, USA). Briefly, Sera diluted(1:10) with sample dilution buffer were mixed with HRP-conjugated RBD solution at a 1:1 volume, and incubated for 30 min at 37°C. The mixture were added into the plate pre-coated with hACE2 receptor, and incubated for 15 min at 37°C. Absorbance was measured on an Epoch Microplate Spectrophotometer (Biotek, VT, USA) at 450 nm. The percentage of inhibition the interaction of RBD and hACE2 was calculated as neutralization (%) = (1-OD value of sera/OD value of negative control) x 100 % (Tan, et al, 2020).

### Plaque reduction neutralization test (PRNT)

PRNT was performed at Clinical Research Center of Masan National Tuberculosis Hospital, an affiliate of Korea Disease Control and Prevention Agenecy (KDCA). All the procedures were conducted in the biosafety level 3 (BL-3) laboratory.

Vero cells were seeded at 8 × 10^5^ cells/well in 6 well plates and cultured overnight. After 2-fold dilution of the serum from 1:10 to 1:10, 240, the diluted serum was mixed 1:1 with 400 Plaque forming unit (PFU)/100 μL of SARS-CoV2 viruses and incubated at 37°C for 1 hour. Vero cells were washed with PBS, and then a serum-virus mixture was added. After incubation for 90 minutes to allow viral infection, the culture medium was removed and washed with PBS. Then, DMEM (Gibco, MA, USA) medium containing 2% FBS and 1.5% Agarose was added, solidified at room temperature for 20 minutes, and cultured at 37° C for 72 hours. Then, after 4% formalin was added and fixed at room temperature for 1 hour, the medium was removed, and the cells were stained with 0.4% crystal violet in 20% methanol solution. After washing three times with distilled water, the plate was completely dried. Plaque number was visually counted and the IC_50_ titer was calculated.

## Discussion

Even in the current situation, we are experiencing the profound health threat of COVID-19 to humanity on a daily basis, hearing news of new infections and unfortunate victims of COVID-19 caused by SARS-CoV-2.

Although several types of COVID-19 vaccines have been developed and used, it is worth noting these three types of vaccines: mRNA vaccine (Moderna, Pfizer-BioNTech, and Curevac), adenovirus vaccine (Oxford-AstraZeneca and Johnson & Johnson) and recombinant protein vaccine (Novavax), and the governments of each country are actively carrying out the vaccination. The current supply of the COVID-19 vaccine is limited compared to demand, and as new variants such as strains B.1.1.7, P.1, B.1.427 / B.1.429 and B.1.351 (Wu et al., 2021) continue to emerge, the effectiveness of existing vaccines being developed against the Wuhan strain of COVID-19 is questioned. Therefore, it is necessary to develop new vaccines to meet the demand for vaccines and ensure efficient vaccines against numerous variants of COVID-19.

Since the mRNA vaccine can be developed in a relatively short time compared to inactivated or recombinant protein vaccines, we initiated the development of an mRNA vaccine against COVID-19 to respond to the urgent demand for vaccines and the future needs of vaccines against emerging variants. In this report, we generated an mRNA vaccine candidate coded for a full-length spike glycoprotein, EG-COVID, European mutant strain (D614G) with 2P-3Q substitution (Bangaru et al., 2020). When we initiated the development of the COVID-19 vaccine, European strains started to emerge and became prevalent, so we predicted that it would be appropriate to develop using the European strain rather than the Wuhan variant. Furthermore, the 2P-3Q substitution is expected to show better efficacy in our study using artificial intelligence analysis and vaccine development research. Currently available vaccines are mostly developed based on the Wuhan strain and all of them have been reported to be less effective against the South African variants (Wu et al., 2021). Our vaccine was developed based on a European variant to which the 2P-3Q substitution was applied and is expected to demonstrate the efficacy against the South African variant. Interestingly, Novavax’s vaccine targeting the same European variant (D614G) with 2P-3Q substitution as our EG-COVID, showed 95.6% effectiveness against the original variants and also for the newer variant (85.6% for the UK variant B.1.1.7 and 60% for the South Africa variant B.1.351) (Mahase, 2021). With this in mind, we anticipated that EG-COVID would have a similar effect against the original variant and new variants as the Novavax’s vaccine.

The human immune system has an effective defense system when attacked by viruses or bacteria. In this sense, IFN-γ plays an important role in the recognition and elimination of pathogens (Kak et al, 2018). We demonstrated that immunization with EG-COVID in mice induced anti-RBD (Receptor Binding Domain) antibodies and SARS-CoV-2 neutralizing antibodies with two doses. Also, we found that EG-COVID induced cellular immune response. Therefore, we concluded that EG-COVID successfully induced virus specific adaptive immune response, which could play an effective antiviral role against SARS-CoV-2. Spike protein and RBDs are the primary targets of neutralizing antibodies because they prevent the virus from binding to airway epithelial cells through invasion of the ACE 2 receptor (Cox et al., 2020). We demonstrated EG-COVID generated neutralizing antibodies against recombinant RBD protein. Since there was an approach whereby neutralizing antibodies directed primarily to RBD of spike protein could be applied as a complementary option to treat severe forms of COVID-19 (Assadiasl et al., 2020), the neutralizing antibody titer induced by EG-COVID could be an important indicator of an effective vaccine against SARS-CoV-2 infection.

All currently available mRNA COVID-19 vaccines are supplied in liquid form and requires special cold chain for storage and transportation of the vaccines. This is a factor that increases the cost of supplying the vaccines and at the same time makes it difficult to distribute the vaccines. This results in the low accessibility of mRNA vaccines, especially for developing countries. To overcome this, it is necessary to prepare the mRNA vaccine in a form that can be stored at room temperature or refrigerated at 4°C, and lyophilization is essential for this purpose. As we observed in our study, EG-COVID is being developed in a form that can be lyophilized and the success of our vaccine will promote equitable access to mRNA vaccines worldwide, as well as establish a system that can deliver vaccines to everyone who needs it in a timely manner.

In conclusion, EG-COVID is a potent and safe vaccine against SARS-CoV-2 that can be stored for a prolonged time in a conventional refrigerator.

## References

1. Assadiasl S, Fatahi Y, Zavvar M, Nicknam MH. COVID-19: Significance of antibodies. Hum Antibodies. 2020;28(4):287–297

2. Bangaru S, Ozorowski G, Turner HL, et al. Structural analysis of full-length SARS-CoV-2 spike protein from an advanced vaccine candidate. Science. 2020;370(6520):1089–1094. doi:10.1126/science.abe1502

3. Corbett K.S., N ENGL J MED Corbett, K. S. et al. SARS-CoV-2 mRNA vaccine design enabled by prototype pathogen preparedness. Nature https://doi.org/10.1038/s41586-020-2622-0 (2020).

4. Cox, R.J., Brokstad, K.A. Not just antibodies: B cells and T cells mediate immunity to COVID-19. Nat Rev Immunol 20, 581–582 (2020).

5. Cunningham AL, Lal H, Kovac M, et al. Efficacy of the herpes zoster subunit vaccine in adults 70 years of age or older. N Engl J Med. 2016;375(11):1019–1032

6. Ickenstein L.M., Garidel P. Lipid-based nanoparticle formulations for small molecules and RNA drugs. Expert Opin. Drug Deliv. 2019;16:1205–1226.

7. Kak G, Raza M, Tiwari BK. Interferon-gamma (IFN-γ): Exploring its implications in infectious diseases. Biomol Concepts. 2018 May 30;9(1):64–79.

8. Karmali PP, Chaudhuri A. Cationic liposomes as non-viral carriers of gene medicines: resolved issues, open questions, and future promises. Med Res Rev. 2007 Sep;27(5):696–722

9. Kim M, Jeong M, Hur S, Cho Y, Park J, Jung H, Seo Y, Woo HA, Nam KT, Lee K, Lee H. Engineered ionizable lipid nanoparticles for targeted delivery of RNA therapeutics into different types of cells in the liver. Sci Adv. 2021 Feb 26;7(9):eabf4398.

10. Lal H, Cunningham AL, Godeaux O, Chlibek R, Diez-Domingo J, Hwang SJ, Levin MJ, McElhaney JE, Poder A, Puig-Barberà J, Vesikari T, Watanabe D, Weckx L, Zahaf T, Heineman TC; ZOE-50 Study Group. Efficacy of an adjuvanted herpes zoster subunit vaccine in older adults. N Engl J Med. 2015 May 28;372(22):2087–96.

11. Lian, T.; Ho, R. J. Trends and developments in liposome drug delivery systems. J. Pharm. Sci. 2001, 90, 667–80.

12. Mahase E. Covid-19: Novavax vaccine efficacy is 86% against UK variant and 60% against South African variant. BMJ. 2021 Feb 1;372:296.

13. Mai Y, Guo J, Zhao Y, Ma S, Hou Y, Yang J. Intranasal delivery of cationic liposome-protamine complex mRNA vaccine elicits effective anti-tumor immunity. Cell Immunol. 2020 Aug;354:104143.

14. Reichmuth, A. M., Oberli, M. A., Jeklenec, A., Langer, R. & Blankschtein, D. mRNA vaccine delivery using lipid nanoparticles. Ther. Deliv. 7, 319–334 (2016).

15. Ryu JI, Park SA, Wui SR, et al. A De-O-acylated Lipooligosaccharide-Based Adjuvant System Promotes Antibody and Th1-Type Immune Responses to H1N1 Pandemic Influenza Vaccine in Mice. Biomed Res Int. 2016;2016:3713656.

16. Sabnis S, Kumarasinghe ES, Salerno T, Mihai C, Ketova T, Senn JJ, Lynn A, Bulychev A, McFadyen I, Chan J, Almarsson Ö, Stanton MG, Benenato KE. A Novel Amino Lipid Series for mRNA Delivery: Improved Endosomal Escape and Sustained Pharmacology and Safety in Non-human Primates. Mol Ther. 2018 Jun 6;26(6):1509–1519. doi: 10.1016/j.ymthe.2018.03.010. Epub 2018 Mar 14.

17. Sercombe L, Veerati T, Moheimani F, Wu SY, Sood AK, Hua S. Advances and Challenges of Liposome Assisted Drug Delivery. Front Pharmacol. 2015 Dec 1;6:286.

18. Tan CW, Chia WN, Qin X, et al. A SARS-CoV-2 surrogate virus neutralization test based on antibody-mediated blockage of ACE2-spike protein-protein interaction. Nat Biotechnol. 2020;38(9):1073–1078

19. Torchilin VP. Recent advances with liposomes as pharmaceutical carriers. Nat Rev Drug Discov. 2005 Feb;4(2):145–60.

20. Vartak A, Sucheck SJ. Recent Advances in Subunit Vaccine Carriers. Vaccines (Basel). 2016 Apr 19;4(2):12. doi: 10.3390/vaccines4020012. PMID: 27104575; PMCID: PMC4931629.

21. Weiss SR, Navas-Martin S. Coronavirus pathogenesis and the emerging pathogen severe acute respiratory syndrome coronavirus. Microbiol Mol Biol Rev. 2005 Dec;69(4):635–64.

22. World Health Organization. “WHO Director-General’s opening remarks at the media briefing on COVID-19—11 March 2020”. Available at: https://www.who.int/dg/speeches/detail/who-director-general-s-opening-remarks-at-the-media-briefing-on-covid-19 11-march-2020. Accessed DEC 1, 2020

23. Wolff JA, Malone RW, Williams P, Chong W, Acsadi G, Jani A, et al. Direct gene transfer into mouse muscle in vivo. Science 1990;247(4949 Pt 1): 1465–8.

24. Wu K, Werner AP, Koch M, Choi A, Narayanan E, Stewart-Jones Gbe, Colpitts T, Bennett H, Boyoglu-Barnum S, Shi W, Moliva JI, Sullivan NJ, Graham BS, Carfi A, Corbett KS, Seder RA, Edwards DK. Serum Neutralizing Activity Elicited by mRNA-1273 Vaccine. N Engl J Med. 2021 Mar 17.

25. Wui SR, Kim KS, Ryu JI, Ko A, Do HTT, Lee YJ, Kim HJ, Lim SJ, Park SA, Cho YJ, Kim CG, Lee NG. Efficient induction of cell-mediated immunity to varicella-zoster virus glycoprotein E co-lyophilized with a cationic liposome-based adjuvant in mice. Vaccine. 2019 Apr 3;37(15):2131–2141.

26. Zhu N, Zhang D, Wang W, et al. A novel coronavirus from patients with pneumonia in China, 2019. N Engl J Med 2020;382(8):727–733.

